# Gut microbial profiling of COVID-19 patients in Uganda

**DOI:** 10.1101/2024.06.28.601197

**Authors:** David Patrick Kateete, Christopher Lubega, Ronald Galiwango, Emmanuel Nasinghe, Monica Mbabazi, Daudi Jjingo, Alison Elliott

## Abstract

**Background:** While COVID-19 spread globally, the role of the gut microbiota in patient outcomes has remained an area of exploration especially in resource limited settings. This study aimed to comprehensively profile the gut microbiome among Ugandan COVID-19 patients and infer potential implications.

**Methods:** Nasopharyngeal swabs, stool, clinical and demographic data were collected from COVID-19 confirmed cases at the COVID-19 isolation and treatment centers in Kampala and Entebbe, Uganda, during the first and second waves of the pandemic in Uganda (i.e., 2020 and 2021, respectively). SARS-CoV-2 presence in the swab samples was confirmed by quantitative real-time RT-PCR assays. 16S rRNA metagenomic next-generation sequencing was performed on the DNA extracted from the stool samples, followed by bioinformatics analysis. Machine learning was used to determine microbes that were associated with disease severity.

**Results:** We observed varied gut microbial composition between COVID-19 patients and healthy controls. Potentially pathogenic bacteria such as Klebsiella oxytoca, Salmonella enterica and Serratia marcescens had an increased presence in COVID-19 disease states, especially severe cases. Enrichment of opportunistic pathogens, such as Enterococcus species, and depletion of beneficial microbes, like Alphaproteobacteria, was observed between mild and severe cases. Machine learning identified age and microbes such as Ruminococcaceae, Bacilli, Enterobacteriales, porphyromonadaceae, and Prevotella copri as predictive of severity.

**Conclusion:** These findings suggest that the microbiome plays a role in the dynamics of SARS-CoV-2 infection in African patients. The shift in abundance of specific microbes can moderately predict severity of COVID-19 in this population. Their direct or indirect roles in determining severity should be investigated further for potential therapeutic interventions.

## Background

Evidence suggests alterations in the composition of gut and other microbiomes including oral and nasopharyngeal following severe acute respiratory syndrome coronavirus 2 (SARS-CoV-2) infection [1–4]. Such alterations could be associated with worse outcomes including increased disease severity and mortality, as well as predisposing patients to co-infections [3,4]. There have been some reports on coinfection in COVID-19 patients, and thus far, the results have varied among different populations [5]. Bacterial co-infections were found to be the dominant co-infection in COVID-19 patients, with *Streptococcus pneumoniae* being the most common followed *by Klebsiella pneumoniae* [6–8]. While distinguishing colonization from infection presents a challenge particularly in the context of COVID-19 [9–11], the prevalence and characteristics of bacterial coinfection in COVID-19 patients is still not well understood.

Metagenomic next-generation sequencing (mNGS) has been shown to increase pathogen diagnostic rates compared to traditional workflows which largely rely on pathogen-specific tests [12]. mNGS also provides valuable information on the composition of the microbiome. When combined with clinical and demographic data it can be used to better understand interactions and dynamics of microbial pathogens and their environments [13]. However, the use of such high-dimension data presents a challenge of lack of validated computational methods for quantitative assessment of the strength of potential microbiome-phenotype associations [14].

Artificial intelligence / machine learning approaches are increasingly being used in metagenomics to explore host-microbiome associations and their relation to development and progression of disease. This is due to their computational superiority in dealing with high dimensional data compared to traditional approaches [15,17]. In this study, we performed a machine learning based analysis of mNGS, demographic and clinical data from COVID-19 patients in Uganda, aimed at predicting / inferring the microbial pathogens associated with COVID-19 disease states.

## Results

### Participants’ characteristics

Stool samples from a total of 105 participants were sequenced (*16S rRNA* sequencing); the paired-end DNA sequences, together with the clinical and/or demographic data were analyzed for microbial profiles. The participants were categorized as ‘mild’ COVID-19 cases (n=42), ‘severe’ COVID-19 cases (n=58), and ‘healthy’ individuals (n=5, i.e., no COVID-19 symptoms and generally in good health), Table 1. Participants in the ‘severe’ category were generally older with a mean age of 45.5 years compared to a mean age of 32.6 years for participants in the ‘mild’ category. While distribution by gender was relatively balanced across categories, majority of the participants were married (55.2%). Further, majority of the ‘mild’ cases reported no comorbidities (76.2%), whereas the ‘severe’ group had a higher proportion of individuals with at least one comorbidity (43.1%), Table 1.

**Table 1:** Participants’ characteristics.

|  | Mild cases<br>(n=42) | Severe cases<br>(n=58) | Healthy controls<br>(n=5) | Total (n=105) |
| --- | --- | --- | --- | --- |
| Age |  |  |  |  |
| Median [Min, Max] | 28.0 [15.0, 88.0] | 46.5 [0, 85.0] | 22.0 [22.0, 61.0] | 35.0 [0, 88.0] |
|  | Freq (%) | Freq (%) | Freq (%) | Freq (%) |
| Missing | 0 (0%) | 0 (0%) | 2 (40.0%) | 2 (1.9%) |
| Sex |  |  |  |  |
| Male | 17 (40.5) | 29 (50.0) | 3 (60.0) | 49 (46.7) |
| Female | 24 (57.1) | 29 (50.0) | 0 (0) | 53 (50.5) |
| Missing | 1 (2.4) | 0 (0) | 2 (40.0) | 3 (2.9) |
| Marital status |  |  |  |  |
| Married | 21 (50.0) | 36 (62.1) | 1 (20.0) | 58 (55.2) |
| Single | 17 (40.5) | 11 (19.0) | 2 (40.0) | 30 (28.6) |
| Separated | 2 (4.8) | 2 (3.4) | 0 (0) | 4 (3.9) |
| Widowed | 1 (2.4) | 2 (3.4) | 0 (0) | 3 (2.9) |
| Missing | 1 (2.4) | 7 (12.1) | 2 (40.0) | 10 (9.5) |
| Education level |  |  |  |  |
| Tertiary | 9 (21.4) | 12 (20.7) | 3 (60.0) | 24 (22.9) |
| None | 1 (2.4) | 5 (8.6) | 0 (0) | 6 (5.7) |
| Primary | 3 (7.1) | 4 (6.9) | 0 (0) | 7 (6.7) |
| Secondary | 24 (57.1) | 11 (19.0) | 0 (0) | 35 (33.3) |
| Missing | 5 (11.9) | 26 (44.8) | 2 (40.0) | 33 (31.4) |
| Comorbidity |  |  |  |  |
| No | 32 (76.2) | 33 (56.9) | 4 (80.0) | 69 (65.7) |
| Yes | 10 (23.8) | 25 (43.1) | 1 (20.0) | 36 (34.3) |

### Composition and relative abundance of taxa

The composition of the gut microbiome exhibited variability across the three categories: mild, severe, and healthy controls. While *Faecalibacterium prausnitzii* was present in both mild and severe groups, it was the most abundant in the healthy controls, followed by *[Eubacterium] biforme*. A few common species, such as *Eggerthella lenta, Collinsella aerofaciens,* and *Parabacteroides distasonis*, were present in the most abundant species across all groups. There was an over representation of some species in the mild cases and severe cases, with an increase in severe cases compared to mild cases. These species included A*kkermansia muciniphila, Prevotella copri and Prevotella stercorea* Conversely, species like *Bacteroides plebeius* and *Eggerthella lenta* were more abundant in the healthy individuals (See Fig 1)

**Fig 1:**
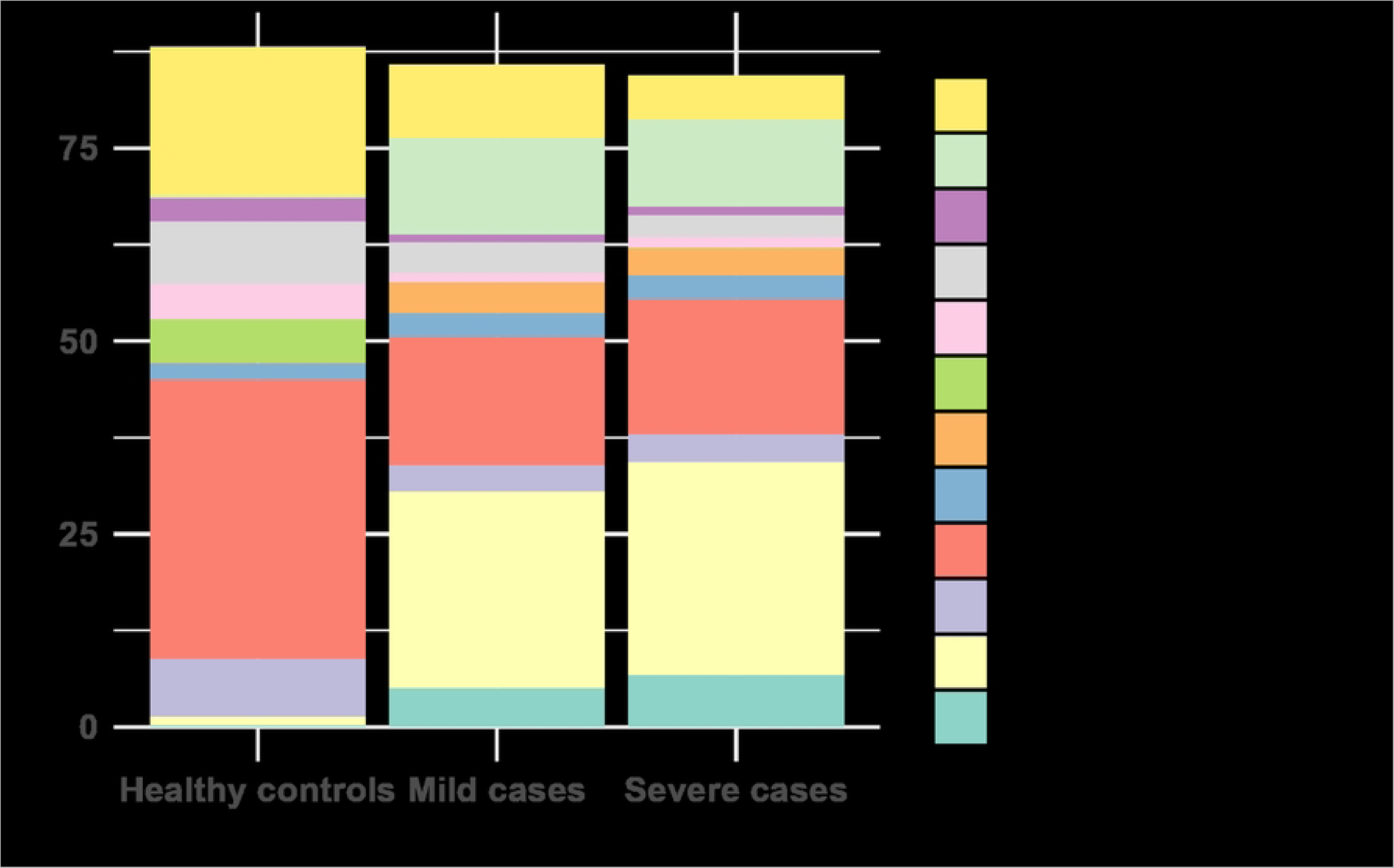
Relative abundance of taxa according to species.

At the genus level, distinctive patterns were observed across the three groups. The genus *Bacteroides* was the most abundant across all groups, however there was a significant reduction among cases compared to healthy controls. There was a significant increase of the *Prevotella*, *Streptococcus*, *Akkermansia* and *Bifidobacterium* genera in cases compared to healthy controls. Similarly, both *Methanobrevibacter* and *Collinsella* were highly abundant in both mild and severe groups while they were less abundant in the healthy controls. *Parabacteroides* and *Ruminococcus* appeared among the most abundant in all groups, but were more highly abundant in the healthy controls. (See Fig. 2).

**Fig 2.**
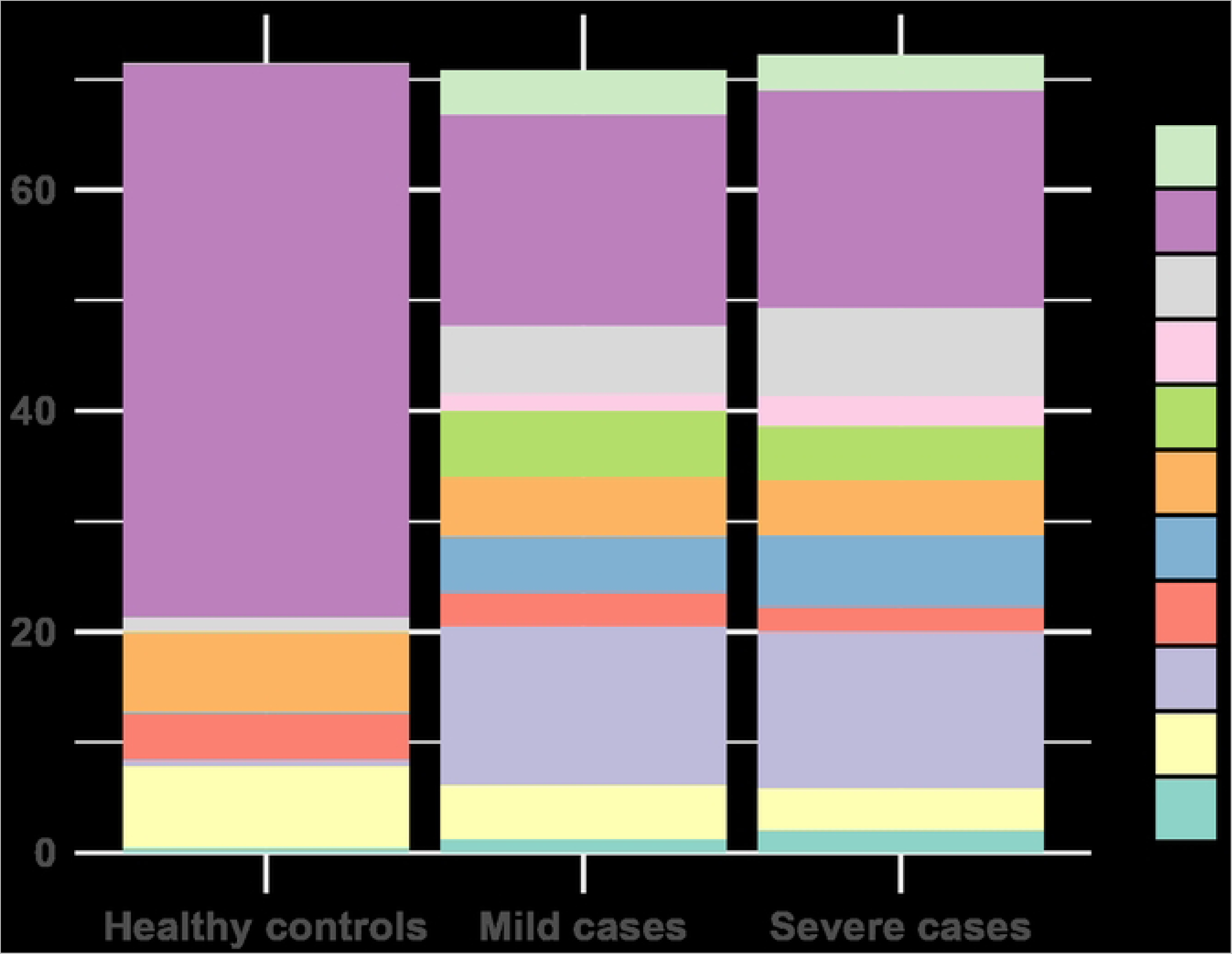
Relative abundance of taxa at genus level.

### Relative abundance of pathogenic species

The relative abundance of species generally considered to be pathogenic was varied across categories. Bacteria such as *Haemophilus influenzae*, *Klebsiella oxytoca*, and *Serratia marcescens* had an increased presence in the disease states, especially severe COVID-19 cases. *Staphylococcus aureus* and *Staphylococcus epidermidis* were both present across the three categories, however, for *S. aureus*, the abundance was highest in severe cases followed by mild cases. *Salmonella enterica* was almost negligible in controls, with a gradual increase in abundance from mild to severe cases. *Klebsiella oxytoca* was absent in the healthy group and had an increasing abundance from mild to severe (Fig 3)

**Fig 3.**
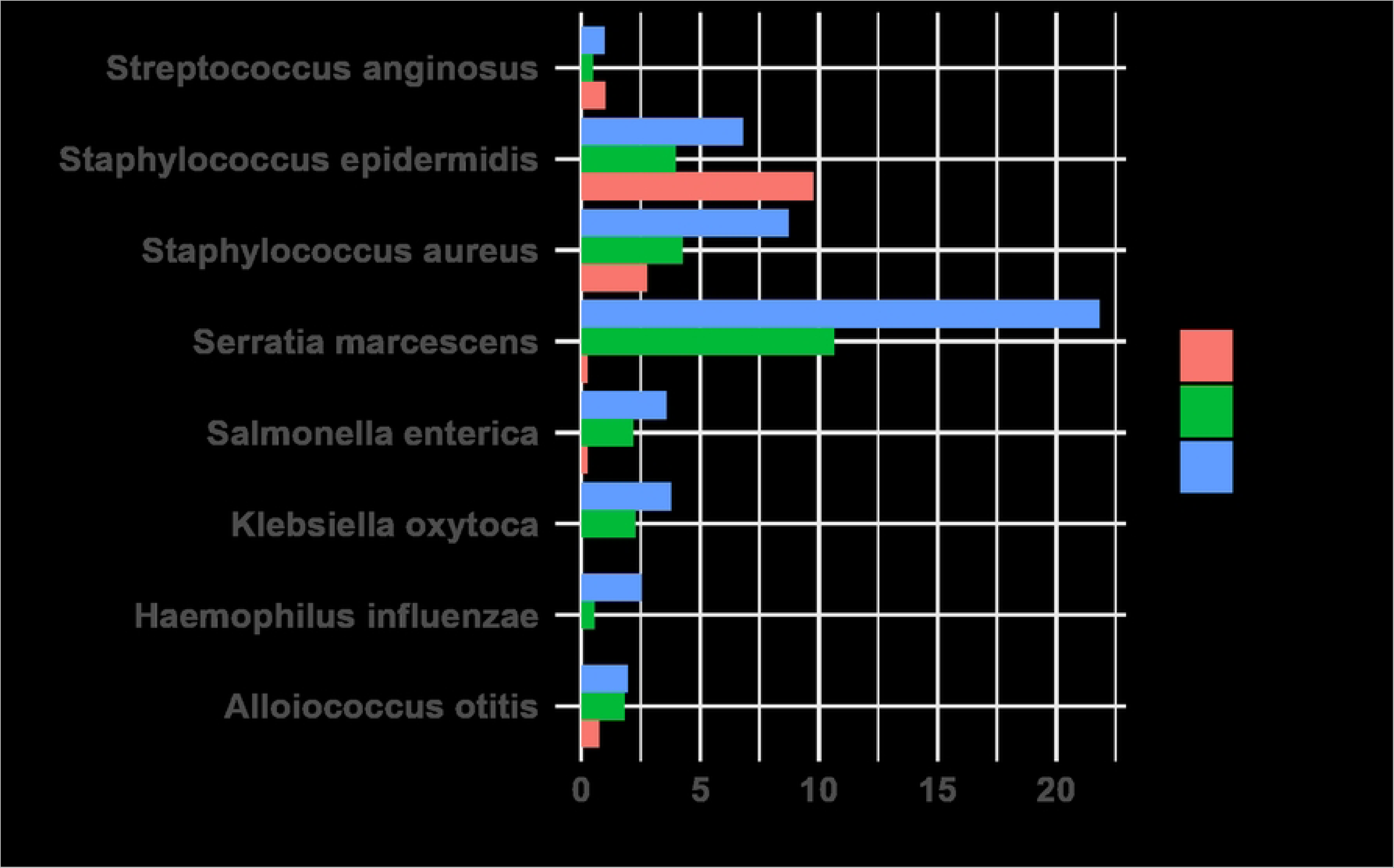
Relative abundance of pathogenic bacteria.

### Diversity of the microbiome

#### Alpha diversity

Investigation of alpha diversity was done using the Shannon diversity index, between the mild and severe cases only, since health controls had only a few samples. The findings suggest comparable microbial diversity between the two groups. Visual inspection of the plots showed no observable differences between the mild and severe cases (See supplementary Fig S1). Further, a statistical analysis using the Wilcoxon rank sum test was consistent with these observations. There was no significant difference in diversity between the two groups (p = 0.223)

#### Beta Diversity

Beta diversity comparison was done using the Bray-Curtis dissimilarity, and Principal Coordinates Analysis (PCoA) was employed to visualize the dissimilarities. The resulting PCoA plot showed three distinct clusters; however, both ‘mild’ and ‘severe’ COVID-19 cases were spread across all three clusters (see supplementary Fig S2). While there was a statistically significant difference in microbial community composition between different severity levels (p=0.013), the severity only explained about 3% of the variance in community composition.

#### Differential abundance analysis

Differential abundance analysis was done using R package DESeq2, to identify significant alterations in the microbial composition across the two levels of COVID-19 severity. Taking mild cases as the reference category, the results showed some OTUs that significantly differ in abundance. *Alphaproteobacteria*, *Anaerostipes*, *Chromatiales*, *Gracilibacteraceae*, *Peptococcaceae*, and *Thermoleophilia* showed a significant decrease in their abundance, with negative log2 fold changes. Among them, *Anaerostipes* displayed the most pronounced decrease with a log2 fold change of-1.69985. Conversely, taxa including *Bacillaceae*, *Bacillales*, *Bacilli*, *Enterococcaceae*, and *Enterococcus* were significantly higher in abundance in severe cases. The *Enterococcaceae* family and the *Enterococcus* genus showcased the steepest increases with log2 fold changes of 2.817143 and 2.869012, respectively (see Table 2)

**Table 2.** Differential abundance of taxa in mild vs. severe COVID-19 cases.

|  | baseMean | log2FoldChange | lfcSE | stat | pvalue | padj |
| --- | --- | --- | --- | --- | --- | --- |
| *Alphaproteobacteria* | 4.101 | -1.58 | 0.398 | -3.97 | 0.016 | 0.007 |
| *Anaerostipes* | 1.431 | -1.7 | 0.381 | -4.463 | 0.005 | 0.002 |
| *Bacillaceae* | 5.875 | 1.924 | 0.473 | 4.069 | 0.016 | 0.007 |
| *Bacillales* | 10.347 | 1.732 | 0.405 | 4.272 | 0.01 | 0.004 |
| *Bacilli* | 180.808 | 1.493 | 0.378 | 3.952 | 0.016 | 0.007 |
| *Chromatiales* | 0.135 | -1.402 | 0.356 | -3.94 | 0.016 | 0.007 |
| *Enterococcaceae* | 7.315 | 2.817 | 0.525 | 5.368 | 0.000 | 0.000 |
| *Enterococcus* | 7.006 | 2.869 | 0.551 | 5.205 | 0.000 | 0.000 |
| *Gracilibacteraceae* | 0.502 | -1.959 | 0.497 | -3.942 | 0.016 | 0.007 |
| *Peptococcaceae* | 3.405 | -1.239 | 0.294 | -4.217 | 0.01 | 0.004 |
| *Thermoleophilia* | 0.06 | -1.618 | 0.412 | -3.924 | 0.016 | 0.007 |

A graphical representation of the differential abundance is shown in Fig 4 where the points are the different OTUs. Blue points represent significant OTUs while grey points represent non-significant OTUs. Below the line are OTUs that decreased among severe cases compared to mild cases.

**Fig 4.**
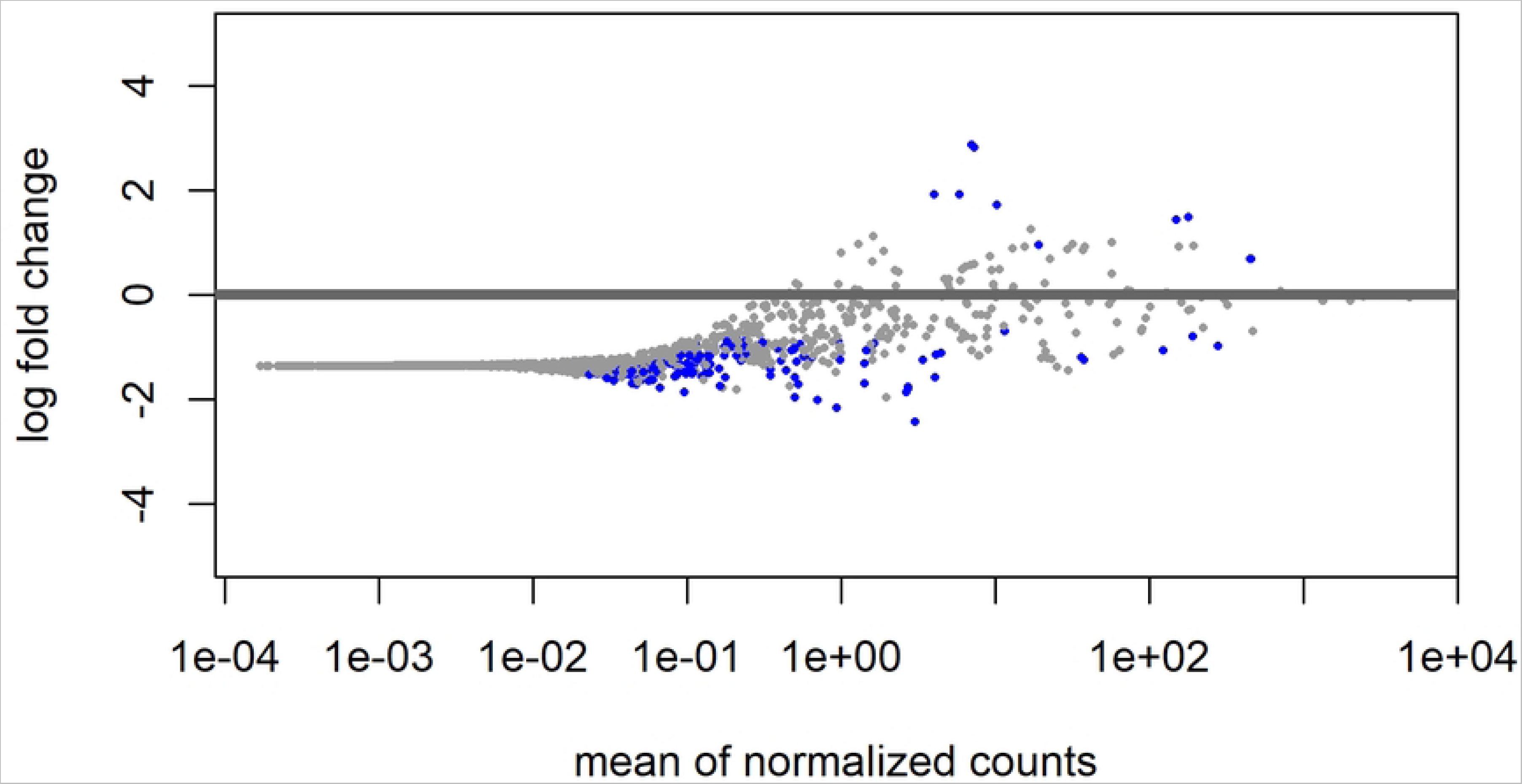
Differential abundance of OTUs in mild vs. severe COVID-19 cases.

### Machine learning prediction of COVID-19 severity

We predicted severity of COVID-19 cases using diverse data points including the demographic characteristics and microbial composition of each sample. Requiring hospitalization was used a proxy for categorizing cases as severe or mild. We included sex, age and presence or absence of comorbidities as the other predictors. A range of models were explored including Logistic Regression which achieved an accuracy of 60.0%, Gradient Boosting which achieved 70.0%, Neural Networks which achieved 70.0%, and the Random Forests model, which was the top performer with an accuracy of 83.3%. Area under curve metrics for the each of the models are shown in supplementary figure Fig S3.

Utilizing predictions from the random forests model, we extracted feature importance metrics and ranked the most important features (taxa and factors) in the prediction of disease severity. Alongside age, several microbial taxa emerged as influential predictors, including *Ruminococcaceae, Bacilli, Enterobacteriales, Porphyromonadaceae, Erysipelotrichales, Bifidobacteriaceae, and Prevotellaceae* (See supplementary Fig S4).

## Discussion

The composition of the gut microbiome in Uganda exhibited variability across the three categories: mild COVID-19, severe COVID-19, and healthy individuals. A few common species, such as *Eggerthella lenta*, and *Parabacteroides distasonis*, were present among the most abundant species across all groups, signifying their prevalent role in the gut ecosystem. On the other hand, some species like *Prevotella copri* and *Akkermansia muciniphila* increased significantly in mild and even higher in severe cases, suggesting their abundance might be influenced by health status.

At the genus level, distinctive patterns were observed across the three groups. The genus *Bacteroides* was consistently present across all three categories, indicating its prevalent role in the gut environment. The increase in presence of *Prevotella*, *Akkermansia, Methanobrevibacter, Bifidobacterium* and *Collinsella* could imply potential interactions or shifts due to the disease state. Overall, while some microbes were shared across the groups, distinct patterns in their abundance offer insights into the microbial dynamics at play in the gut. Similar studies in other populations have reported changes in the composition of the gut microbiota during diseased states [1,2,18–20]. The increase or decrease in abundance of specific microbes could provide hints about their roles in health and disease, and further investigations into their functionalities might shed light on their significance in the context of COVID-19.

Decreased microbial diversity correlated with higher disease severity. This aligns with existing literature which has emphasized the protective role of microbial diversity against pathogenic challenges. Promoting a diverse microbiome through dietary and therapeutic interventions could potentially reduce the severity of diseases like COVID-19 [21,22]. Other studies in other populations have observed correlations between microbial composition and viral replication efficiency [22,23]. Understanding which microbes facilitate or hinder viral replication can lead to targeted therapeutic approaches.

Enrichment of opportunistic pathogens, such as *Enterococcus* species, and depletion of beneficial microbes, like *Alphaproteobacteria*, was observed in mild and severe COVID-19 cases. A review of previous research [24] suggested similar findings, noting a decrease in beneficial gut microbes with anti-inflammatory properties in COVID-19 patients. The increase in opportunistic pathogens is consistent with reports that COVID-19 patients frequently experience secondary bacterial infections, which can worsen disease outcomes [25–27].

Previous studies incorporated machine learning to predict disease severity based on microbial compositions [28,29]. Distinct microbial signatures were identified, with the prevalence of specific OTUs in COVID-19 patients and a particular abundance of *Prevotella* in severe cases. This was similar to findings from a recent study [30] which highlighted a protective role of the *Prevotella* genus in the long-term recovery process while other investigations [31,32] reported varied results. The variation could be attributed to various factors like sample collection methods, patient demographics, or even geographic variations in microbial populations. If *Prevotella* or other bacteria play a direct or indirect role in disease exacerbation, interventions targeting these bacteria might be explored as a treatment or preventive strategy.

### Limitations

We had several limitations in our study including the imbalance between cases and healthy controls. The ongoing lockdown during the enrollment period made it difficult to enroll healthy individuals due to movement restrictions. As such, statistical comparisons between cases and health controls were not included in the results. Additionally, some factors like prior antibiotic usage and diet which might influence the microbiome, were not elaborately reported and investigated. We also used to require hospitalization as a practical proxy for symptom severity in our study in place of direct severity measures, which can be subjective.

In conclusion, it is evident that the microbiome plays a significant role in SARS-CoV-2 infection dynamics in African patients. The enrichment of opportunistic pathogens and depletion of beneficial microbes in COVID-19 patients align with findings from prior studies emphasizing the potential therapeutic implications of such microbial shifts. Specifically, an elevated abundance of bacteria like *Prevotella* and *Akkermansia* may offer insights into their role in disease severity. Additionally, reduced microbial diversity in severe cases is suggestive of a protective effect of a diverse microbiome as reported in recent studies [19–21].

Given the evident associations between specific taxa and COVID-19 severity, more comprehensive and longitudinal studies are necessary. This will help in understanding causative relationships from mere correlations, which is vital for therapeutic applications. The utility of machine learning models in predicting disease severity based on microbial compositions suggests a promising avenue for precision medicine.

## Materials and methods

### Study design

The study period was June 2020 to December 2022. We used a case-control study nested in a parent study in which we conducted 16S metagenomic sequencing of stool samples of 100 confirmed COVID-19 patients and 5 healthy controls. Under the parent study titled “Establishment of Quality Assured COVID-19 Specimen Repository to Support Research in Diagnosis, Prevention and Management of SARS CoV2 in Uganda”, samples were collected alongside demographic and clinical data during the COVID-19 pandemic in Uganda.

### Data collection and storage

Samples were collected based on the clinical COVID-19 suspicion of individuals who presented with symptoms of acute respiratory infection. For each individual, a stool sample was collected using sterile cotton swabs. Sample collection took place at Mulago hospital, other ministry of health approved COVID-|9 treatment centres and the community. The samples were processed in the Makerere University Laboratories and stored at the Integrated Biorepository of H3Africa Uganda [35]. The COVID-19 diagnosis was performed by SARS-CoV-2 detection using real-time reverse transcriptase-polymerase chain reaction (RT-PCR) assay on the gut swabs. The patients were categorized into two groups based on disease severity, i.e., mild, and severe cases.

Healthy controls were individuals recruited from the general population with no illness and antibiotic use in the last 3 months, and tested negative for SARS-CoV-2. All the enrolled subjects signed informed consent to have their samples stored for research purposes. An aliquot of the swabs was stored at-80°C.

Data including demographics, laboratory results and medical therapy of all individuals was collected during specimen collection using an approved study specific case report form and stored in an electronic database. We used de-identified data in all analyses.in accordance with good clinical practice principles.

### Metagenomic Next-generation Sequencing

#### Sample Retrieval

Stool samples were retrieved from the repository with strict adherence to protocols to guarantee their viability. Details such as the storage date, temperature consistency, and any exposure to freeze-thaw cycles were cross-checked. During retrieval, samples remained shielded in dry ice to preserve their integrity. The viability of the samples was ascertained by assessing the quality of microbial DNA. Samples which exhibited signs of degradation, contamination, or which met the exclusion criteria described above were earmarked as non-viable and excluded.

#### Microbial DNA Extraction

Microbial DNA was efficiently extracted from each Stool sample using the TIANamp Micro DNA Kit (DP316, TIANGEN BIOTECH). This procedure was done in adherence to the manufacturer’s guidelines, ensuring high yield and purity of the DNA for downstream processes.

#### Sequencing

The single-stranded DNA fragments were captured on a flow cell where they underwent bridge amplification, forming clusters. Each cluster was then sequenced on the Illumina MiSeq platform [36] from both ends i.e., paired end sequencing to obtain the raw reads. The paired-end reads were demultiplexed based on their unique indices.

### Bioinformatics Analysis

Taxonomic Classification was done using Kraken 2, which uses a k-mer based approach to compare fragments of the metagenomic sequences to a user-specified database, enabling the classification of the microbial composition of each sample [37]. For this analysis the Greengenes database, a well-curated reference database specializing in 16S ribosomal RNA sequences was used [38]. Subsequent analyses were done using R Vegan [39], Phyloseq [40] and DESeq2 [41] packages.

### Statistical Analysis

All plots and statistical analyses were conducted with R v4.3.1 [42], and the vegan package in R was used to obtain the diversity indexes, including the Shannon index for alpha diversity and Bray–Curtis dissimilarities for beta diversity. The differences in alpha diversity among or between groups were statistically evaluated using Kruskal-Wallis test. The statistical differences between these beta-diversity indices were computed using permutational multivariate analysis of variance (PERMANOVA) with the adonis() function in the vegan package [39]. Differential abundance analysis was done using the DESeq2 package [41] and P-values were adjusted for multiple testing using the false-discovery rate correction.

### Machine Learning

We predicted severity of COVID-19 cases using demographic characteristics and microbial composition of each sample. Following pre-processing, the data was split into training and testing sets. Several models were trained including Logistic Regression, Random Forests, Gradient Boosting, and Neural network followed by computation of accuracy metrics to evaluate model performance. The best performing model was used to predict clinical severity of COVID-19 from microbial profiles. Feature importance metrics were used to obtain taxa that are significantly associated with disease severity.

### Ethical Considerations

Ethical approval for this study was sought from the Makerere University School of Biomedical Sciences Research Ethics Committee (SBS-REC) to conduct this study (approval # SBS-2023-330). The CoVBank project in which this study was nested also obtained ethical approvals and written informed consent from all the study participants. The SBS-REC waived the need for re-consenting the participants as the prior written consent included consent for storage of samples and future use of the stored samples. All patient data and samples were deidentified in accordance with good clinical practice principles. IBRH3AU has ethics approval from the SBS-REC and from the Uganda National Council for Science and Technology (UNCST) to collect, process, store, and share biospecimens. Further still, IBRH3AU obtained ethics approval from the Mulago National Hospital REC and UNCST to collect biospecimens from all COVID-19 cases receiving care from various treatment centers under the Uganda’s Ministry of Health. Participants consented to sample storage and use of their samples and data in current and future studies related to SARS-CoV-2 infection.

## Acknowledgements

This work was supported by a grant from the Science for Africa Foundation Reference # GCA/SARSCov2-2-20-006 (to AE). The authors extend their appreciation to the COVBANK project investigator for granting access to samples and data that was used in this work.

## Supplementary Material

### Supplementary Figures

S1 Alpha diveristy between mild and severe cases

S2 Beta diveristy between mild and severe cases

S3 Area under the curve metrics for machine learning models

S4 Feature importance metrics

